# Reconstructed protein sequence evolution consistent with the evolution of C_4_ photosynthesis via a C_2_ ancestor in the Paniceae

**DOI:** 10.1101/733493

**Authors:** Daniel S. Carvalho, Sunil Kumar Kenchanmane Raju, Yang Zhang, James C. Schnable

## Abstract

The grass tribe Paniceae includes a monophyletic subclade of species, the MPC clade, which specialize in each of the three primary C_4_ sub-pathways NADP-ME, NAD-ME and PCK. The evolutionary history of C_4_ photosynthesis in this subclade remains ambiguous. Leveraging newly sequenced grass genomes and syntenic orthology data, we estimated rates of protein sequence evolution on ancestral branches for both core enzymes shared across different C_4_ sub-pathways and enzymes specific to C_4_ sub-pathways. While core enzymes show elevated rates of protein sequence evolution in ancestral branches consistent with a transition from C_3_ to C_4_ photosynthesis in the ancestor for this clade, no subtype specific enzymes showed similar patterns. At least one protein involved in photorespiration also showed elevated rates of protein sequence evolution in the ancestral branch. The set of core C_4_ enzymes examined here combined with the photorespiratory pathway are necessary for the C_2_ photosynthetic cycle, a previously proposed intermediate between C_3_ and C_4_ photosynthesis. The patterns reported here are consistent with, but not conclusive proof that, C_4_ photosynthesis in the MPC clade of the Paniceae evolved via a C_2_ intermediate.

## Introduction

C_4_ plants are responsible for over a quarter of world terrestrial photosynthetic productivity (Gillon and Yakir, 2001; Sage, 2001). C_4_ grasses alone account for approximately 18% of global productivity (Ehleringer et al., 1997; Wand et al., 1999). The high productivity of C_4_ plants is linked to their ability to increase the ratio of CO_2_ to O_2_ around RuBisCO, lowering the net energy negative process of photorespiration (Bull, 1969; Keys, 1986), and reducing water losses to transpiration. C_4_ plants are able to keep their stomata closed for longer because they are less sensitive to declines in intraleaf CO_2_ concentrations than plants dependent on C_3_ photosynthesis. C_4_ photosynthesis is thought to have arisen in parallel in over 60 lineages approximately 30 million years ago in responses to a drop in CO_2_ levels (Vicentini et al., 2008; Sage et al., 2011).

While exceptions exist (Akhani et al., 2003; Offermann et al., 2015), many lineages utilizing C_4_ photosynthesis split the process between two different cell types: mesophyll (M) and bundle-sheath (BS) cells. In the M cells, CO_2_ is fixed into bicarbonate to form oxaloacetate (OAA). OAA is converted to either malate or aspartate or both depending on the species and the C_4_ pathway being used. Malate or aspartate is then transported to the BS cells where it is decarboxylated, releasing CO_2_ which is then fixed via the conventional Calvin-Benson cycle (Kanai and Edwards, 1999). While this broad pattern is consistently found in a wide range of C_4_ using species, the C_4_ photosynthesis cycle can be divided into three subtypes based on 1) the decarboxylase enzyme which releases CO_2_ within the bundle sheath and 2) the linked property of where decarboxylation occurs. The three enzyme families known to act as decarboxylases for C_4_ as NAD-malic enzyme (NAD-ME) which decarboxylates in the mitochondria, NADP-malic enzyme (NADP-ME) which decarboxylates in the chloroplast and PEP carboxykinase (PCK) which decarboxylates in the cytosol (Kanai and Edwards, 1999; Furbank, 2011). Individual species might utilize a single decarboxylase and hence a single C_4_ pathway or multiple decarboxylases and multiple C_4_ pathways (Walker et al., 1997; John et al., 2014; Huang et al., 2016; de Oliveira Dal’Molin et al., 2016; Washburn et al., 2017).

The tribe Paniceae within the grasses includes 84 genera (Morrone et al., 2012). Notably the clade encompasses different species which utilize each of the the three C_4_ pathways as their primary carbon fixation mechanism. These species share a single common ancestor to the exclusion of any know C_3_ lineage (Sage et al., 2011; Washburn et al., 2015).

This clade containing only C_4_ photosynthesizers encompasses the subtribes Melinidinae (PCK subtype), Panicinae (NAD-ME subtype) and Cenchrinae (NADP-ME subtype) and is also referred to as the MPC clade. Ancestral state reconstruction based on phylogenies and current state data identified either NAD-ME subtype of C_4_ photosynthesis or C_3_ photosynthesis as the likely ancestral state of the MPC clade (Washburn et al., 2015). Ancestral state reconstruction based on expression data suggested that the common ancestor may have been a species using all three C_4_ pathways simultaneously (Washburn et al., 2017).

Here we sought to employ a third approach based on the reconstruction of rates of protein sequence evolution along ancestral branches of the phylogeny. Estimates of changes in protein sequence evolution rates between C_3_ lineages and ancestral branches along the phylogeny of the MPC clade were used to evaluate several potential models for the photosynthetic state of the common ancestor of the MPC clade. While several core enzymes shared by all three C_4_ pathways did indeed show elevated rates of protein sequence evolution in the ancestral branch leading to the MPC clade, none of the three decarboxylases showed elevated rates of protein sequence evolution along this branch. At least one gene involved in photorespiration also showed elevated rates of protein sequence evolution in the ancestral MPC branch. Our findings are suggestive of a C_2_ pathway intermediate ancestor at the base of the MPC clade. The C_2_ uses the photorespiratory pathway as a CO_2_ carbon pump and has been proposed as a potential intermediate state between the conventional C_3_ and C_4_ photosynthetic pathways (Tolbert, 1997; Mallmann et al., 2014; Edwards, 2019). Taken together, these results may represent a “ghost of C_2_ past” in the genomes of this C_4_ clade Edwards (2019).

## Material and methods

### Plant growth and RNA-Seq data generation for *Urochloa fusca*

*Urochloa fusca* seeds were planted and grown in a Percival (Percival model E-41L2) growth chamber with target conditions of 111 *µ*mol m^-2^ s^-1^, 60% relative humidity, a 12 hour/12 hour day night cycle with a target temperature of 29°C during the day and 23°C at night. Under the growing conditions employed, twelve days after planting (DAP) plants were collected. The whole seedlings (except root) were used for RNA extraction. RNA isolation and library construction followed the protocol described by Zhang *et al.* (Zhang et al., 2015). The library was sequenced using 2×100 bp paired end Illumina sequencing, and transcripts were generated using Trinity (v2.0.6) with default parameters with the exception of “–seqType fq –max_memory 20G –CPU 16” (Grabherr et al., 2011).

### Sequence data set

Coding Sequences (CDS) for the transcript annotated as “primary” for each gene in *Brachypodium distachyon* (Initiative et al., 2010), *Oryza sativa* (rice) (Kawahara et al., 2013), *Panicum hallii* (Lovell et al., 2018) and *Setaria italica* (foxtail millet) (Bennetzen et al., 2012) were obtained from Phytozome 12 (https://phytozome.jgi.doe.gov/pz/portal.html). For *Dichanthelium oligosanthes* the CDS were retrieved from version v1.001 in CoGe OrganismView, genome ID 28856 (https://genomevolution.org/CoGe/OrganismView.pl) (Studer et al., 2016). CDS from *Pennisetum glaucum* (pearl millet) were obtained from (Varshney et al., 2017). CDS from *Panicum miliaceum* (proso millet) were downloaded from CoGe (Genome ID: 52484), version v1 (Lyons and Freeling, 2008; Zou et al., 2019). Open reading frames for *Urochloa fusca* were obtained from the transcriptome assembly described above. As a number of transcript assemblies were not full length in manual proofing, open reading frames were constrained to those ending in a stop codon, but were not required to start with an ATG.

### Orthology assignment

LASTZ (Harris, 2007) was used to perform all by all comparisons of coding sequence from the primary transcript of each gene as downloaded from from Phytozome 12 with the following parameters –identity=70 –coverage=50 –ambiguous=iupac, –notransition, and –seed=match12. LASTZ output was parsed to identify syntenic orthologs using QuotaAlign with the additional parameters –tandemNmax=10, cscore=0.5, –merge and –Dm=20 (Tang et al., 2011). To reverse the collapse of tandem gene clusters which are part of the QuotaAlign algorithm, final syntenic orthologs were assigned based on the gene copy with the highest LASTZ alignment score within 20 genes up or downstream of the original syntenic location predicted by quota align. Synteny analysis of proso and pearl millet could not retrieve all C_4_ related genes possibly due to larger amounts of genomic rearrangement and smaller blocks of conserved synteny relative to other grass species included in this study. When no syntenic ortholog could be identified in pearl millet and/or proso millet based on synteny, the best reciprocal LASTZ hit to the foxtail millet gene copy was included as a presumed ortholog, even in the absence of synteny. A similar process was employed for four genes from the *Dichanthelium oligosanthes* genome, as this genome was assembled using purely short read technology and remains fragmented into a large number of scaffolds. In all cases, orthology for *Urochloa fusca* sequences was inferred based on reciprocal best LASTZ hits with the foxtail millet C_4_ genes. The list of syntenic orthologous and presumed orthologous gene sequences analyzed here is provided in supplemental material 2.

### dN/dS calculation and evolutionary analyses

The DNA sequence of the coding sequence region of each gene was translated to an amino acid sequence. Amino acid sequences for groups of orthologous and presumed orthologous genes were aligned using Kalign version 2.04 (Lassmann and Sonnhammer, 2005). This amino acid multiple sequence alignment was used to create a codon level DNA sequence alignment. The codon alignment and a guide phylogenetic tree created based on the known relationships of all species included in the analysis (Edwards et al., 2011) were used to calculate the nonsynonymous and synonymous substitution rates (dN/dS) for each branch of the tree using codeml from the PAML package version 4.09 (Yang, 2007).

### Statistical comparison of branch dN/dS values and photosynthetic trait values

A Fisher Exact Test was performed in order to test whether significant differences in evolutionary rate existed between between branches leading to species utilizing C_3_ photosynthesis and branches leading to species utilizing C_4_ photosynthesis. For each comparison, a contingency table was constructed comparing the separate estimated number of synonymous and nonsynonymous substitutions in two branches or groups of branches. For each gene, each one of the C_4_ branches was tested for an elevated ratio of nonsynonymous to synonymous substitutions relative to the aggregate background C_3_ branches. C_3_ background values for both non-synonymous and synonymous substitutions were summed across three branches each leading to a single C_3_ species: *D. oligosanthes, B. distachyon*, and *O. sativa*. A Fisher Exact Test was used to compare each tested branch to the aggregate C_3_ branch values.

## Results

The evolutionary patterns of ten genes encoding enzymes either unique to individual pathways of C_4_ photosynthesis or shared by two or more pathways were investigated (Figure 2). The selection of genes for investigation was guided both by the strength of literature supporting their specific roles in one or more pathways of C_4_ photosynthesis and the identification of high confidence orthologs in as many of the grass species included in this analysis (Table 1). Two different signatures were possible for genes which experienced positive selection as part of a transition from C_3_ to C_4_ photosynthesis. The first of these two potential signatures is an unambiguous signal where the rate of nonsynonymous substitutions (dN) exceeds the rate of synonymous substitutions (dS) on a given branch (i.e. dN/dS >1) (Figure 3). This signal can be observed either on extremely short branches, or when positive selection continued for long periods of time. The second potential signal is more ambiguous. When an interval of positive selection occurs on a given branch but is preceded and/or followed by purifying within the same branch, the effects of these two intervals cannot be disambiguated without including additional outgroups to break up the branch into short segments. As a result, in branches that include both positive and purifying selection dN/dS will be elevated relative to branches experiencing purely purifying selection. However, despite the presence of positive selection, the dN/dS ratio for the entire branch may still still be less than 1 (Figure 3). Unfortunately, the same signature (elevated dN/dS ratios relative to other branches but not exceeding a ratio of one) is also produced in cases of relaxed selection of pseudogenization. Here we considered either case to be potentially informative about the origins of C_4_ photosynthesis in the MPC clade, however the interpretation of all results presented here must be in the context of both potential explanations for elevated substitution rates.

**Table 1:**
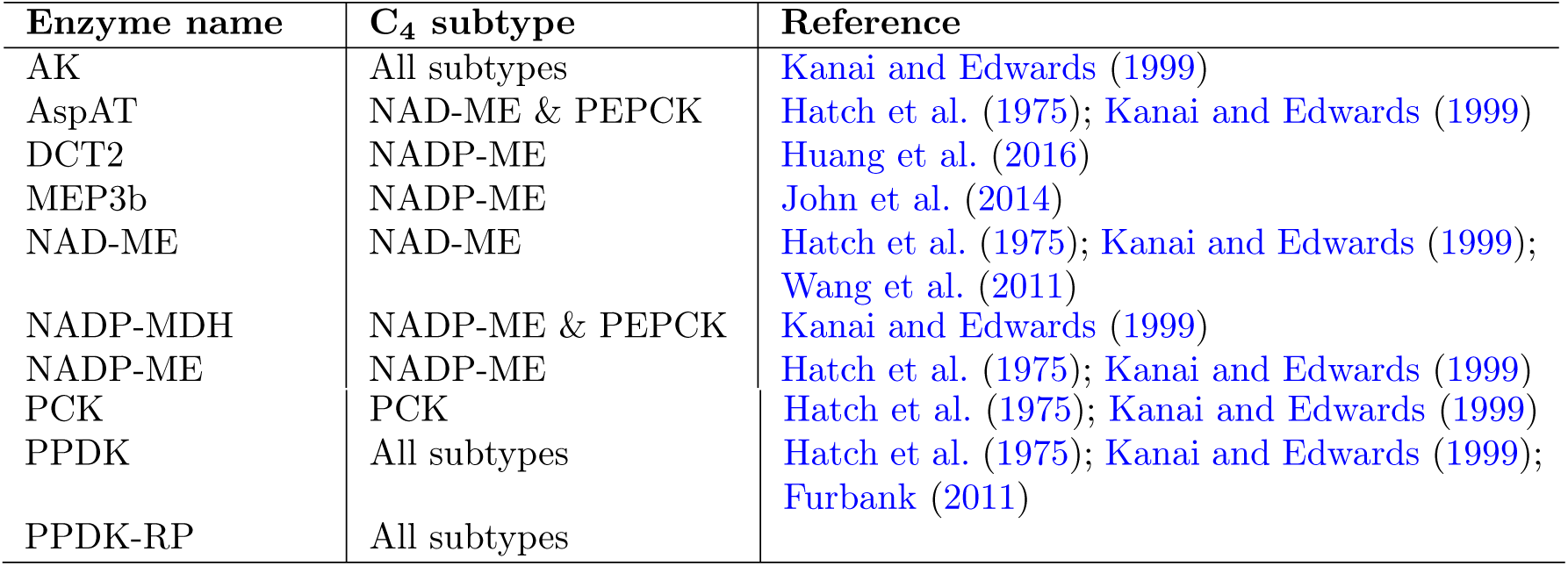
List of enzymes investigated as part of this study, and the set of C_4_ subpathways each enzyme was inferred to contribute to based on the literature.

|  |  |  |
| --- | --- | --- |
| PCK | PCK | Hatch et al. (1975); Kanai and Edwards (1999) |
| PPDK | All subtypes | Hatch et al. (1975); Kanai and Edwards (1999);<br>Furbank (2011) |
| PPDK-RP | All subtypes |  |

**Figure 1:**
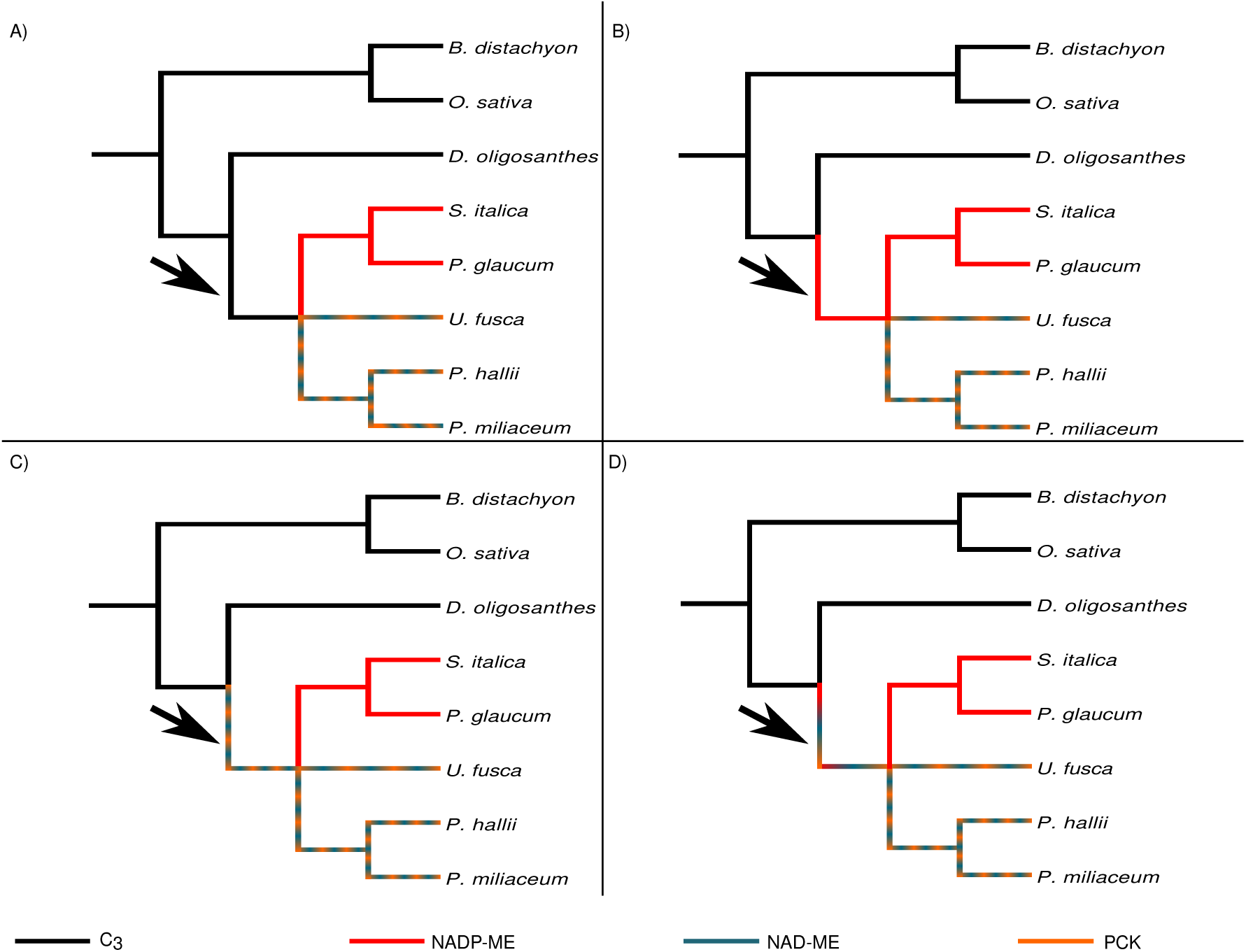
Potential models for the evolution of C_4_ photosynthesis in the MPC clade. A) The common ancestor of the MPC clade utilized C_3_ or another non-C_4_ pathway and C_4_ evolved independently in the three subtribes utilizing different C_4_ pathways Washburn et al. (2015); B) The common ancestor of the MPC clade utilized the C_4_ NADP-ME pathway and the NAD-ME and PCK clades represent later evolutionary changes from one C_4_ subtype to another; C) The common ancestor of the MPC clade utilized either the C_4_ NAD-ME and PCK pathways or a mix of both; D) The common ancestor of the MPC clade utilized utilized all three pathways simultaneously (Washburn et al., 2017). Black arrow indicates the ancestral branch for the MPC subclade within the Paniceae.

**Figure 2:**
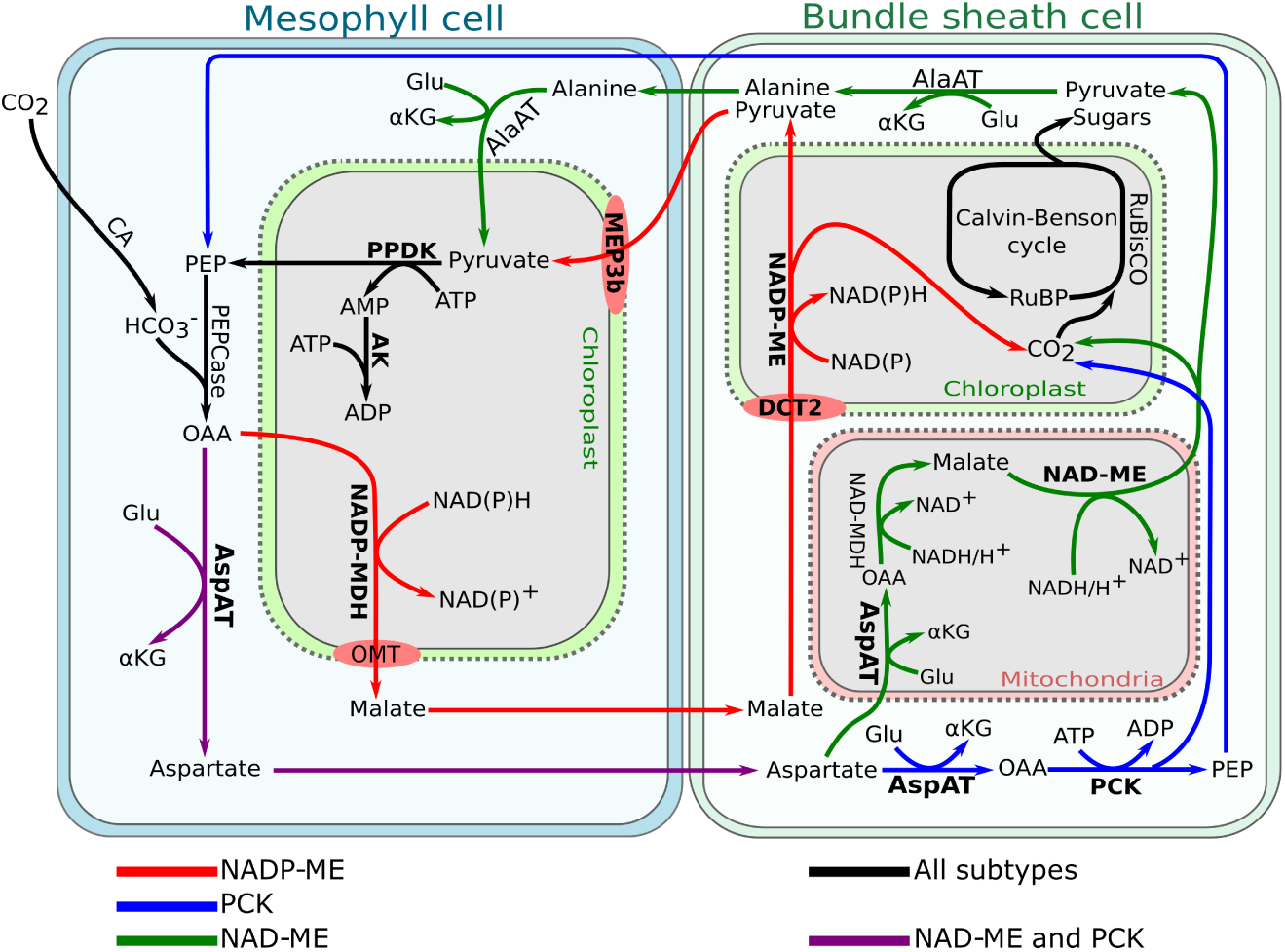
Simplified pathway representation of the three main C_4_ photosynthesis subtypes. Enzymes where protein sequence evolutionary rates were calculated as part of this study are indicated in bold.

**Figure 3:**
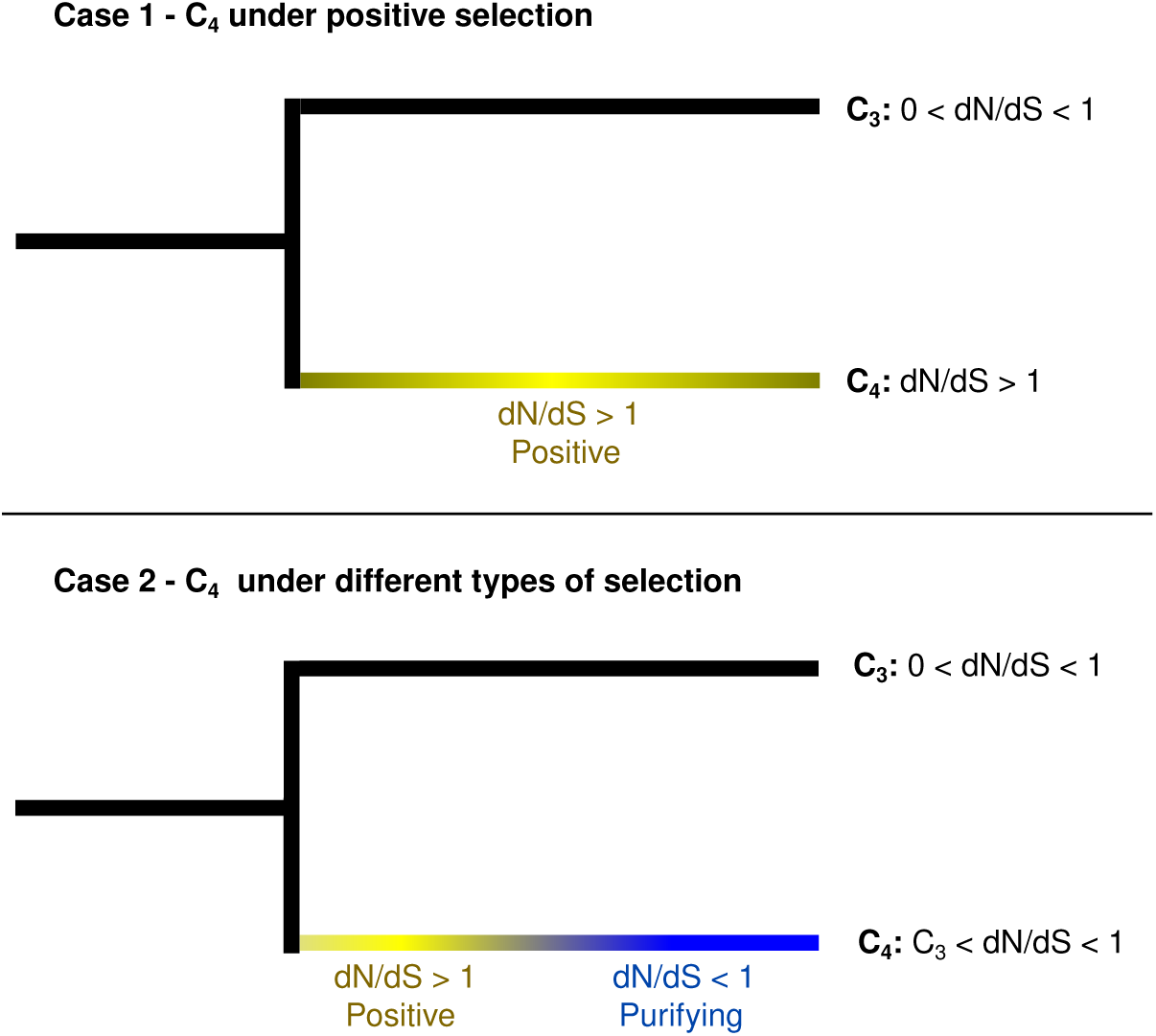
Models for different histories of selection and the predicted outcomes on dN/dS ratios. Case 1 shows a classical case of positive selection leading to change in function, while Case 2 shows a case where an enzyme might have gone through a mixture of positive and purifying selection leading to a change in function and a final purifying selection period to maintain the new enzymatic function.

Core C_4_ enzymes, those which are utilized by all three C_4_ photosynthetic subtypes, showed distinct patterns of change in synonymous/nonsynonymous substitution rates relative to enzymes used in specific subtypes. The analysis of the phylogenetic tree was performed on the branches leading to C_4_ species as well as the ancestor branch of the Paniceae, which is the common ancestor of all C_4_ species in this study (Figure 1). Both PPDK and PPDK-RP, core enzymes, showed a significantly faster rates of protein sequence evolution in the common ancestor of C_4_ species branch relative to C_3_. Adenylate kinase (AK) did not show accelerated protein sequence evolution in the ancestral branch, but showed significantly faster rates on the proso and pearl millet branches compared to their C_3_ counterparts (Figure S1).

None of the subtype specific enzymes showed significantly higher dN/dS values than the background C_3_ genes in the common ancestor of C_4_ species branch (Figure S2). Dicarboxylic acid transporter 2 (DCT2), NADP-ME and MEP3b are all employed in the NADP-ME C_4_ subtype. Both NADP-ME and MEP3b enzymes showed a significantly faster evolutionary rate in all branches of the NADP-ME subtype (foxtail millet, pearl millet and their ancestral branch) and most other C_4_ branches compared to the background C_3_ rate. The C_4_ branches showing no significant differences in dN/dS relative to the background C_3_ rate were: *P. hallii* for the NADP-ME enzyme, and both *P. hallii* and NAD-ME ancestor for MEP3b. DCT2 exhibited a similar pattern to the other two enzymes, with the exception of the branch leading to *S. italica*. An evolutionary pattern shared among NADP-ME, MEP3b and DCT2 enzymes was a significantly faster evolutionary rate in *U. fusca* and proso millet, which perform PCK and NAD-ME subtypes, respectively, compared to the background C_3_ rate. NADP-MDH only showed significantly faster evolutionary rates than C_3_ branches in pearl millet, NADP-ME species, and the ancestral branch of NAD-ME subtype. Both AspAT and NAD-ME showed significantly higher dN/dS ratios in branches leading to NAD-ME and PCK subtype species compared with C_3_ species (Table 2).

**Table 2:**
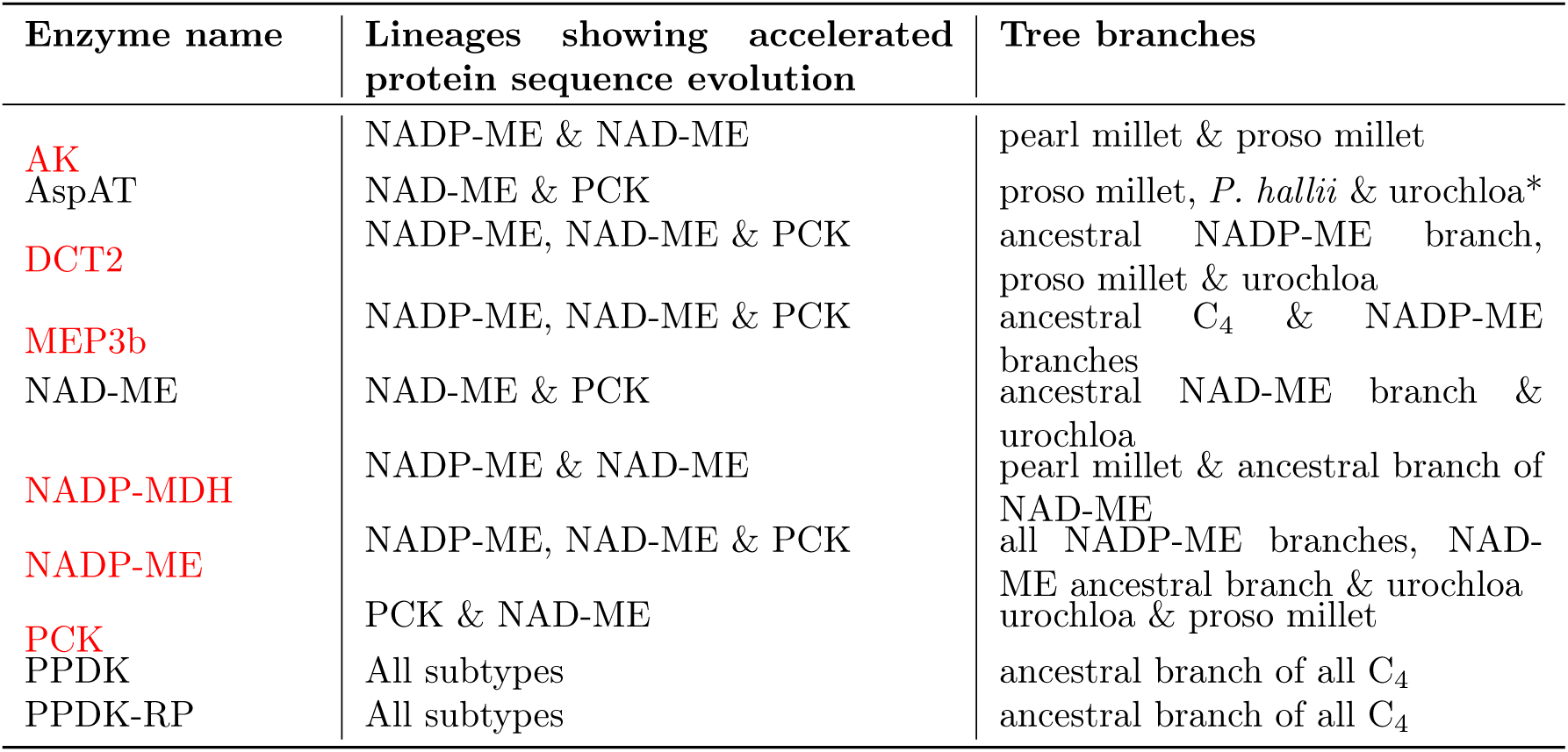
List of enzymes employed in this study which showed elevated rates of protein sequence evolution relative to the C_3_ background in one or more branches leading to C_4_ species. Cases highlighted in red showed elevated rates of protein sequence evolution in at least one branch which is inconsistent with the canonical assignment of MPC grasses into three clades each utilizing a single distinct C_4_ pathway. *this branch was marginally insignificant p = 0.062.

In addition to the 10 C_4_ photosynthesis related enzymes studied here, the approach used to find genes involved in the C_4_ cycle was used to retrieve the sequences of genes encoding two enzymes involved in the photorespiratory pathway: glycolate oxidase (GOX) and serine hydroxymethyl-transferase (SHMT). Both enzymes exhibit faster rates of protein sequence change in branches leading to C_4_ species than in branches leading to C_3_ species. GOX is localized in the peroxisome while SHMT is localized in the mitochondria (Figure 4). The evolutionary pattern of these enzymes were different. GOX showed a faster evolutionary rate than C_3_ species branches in the ancestral branch of all Paniceae, the branch leading to *U. fusca*, the ancestral branch of *S. italica* and *P. glaucum* and the branch leading to *S. italica*. On the other hand, SHMT showed fast evolving branches in *U. fusca* branch, the ancestral branch of both *P. miliaceum* genes and one of the *P. miliaceum* genes. In the branch leading to the common ancestor of the MPC clade, GOX, but not SHMT, also showed elevated rates of protein sequence changes relative to branches leading to C_3_ species S3).

**Figure 4:**
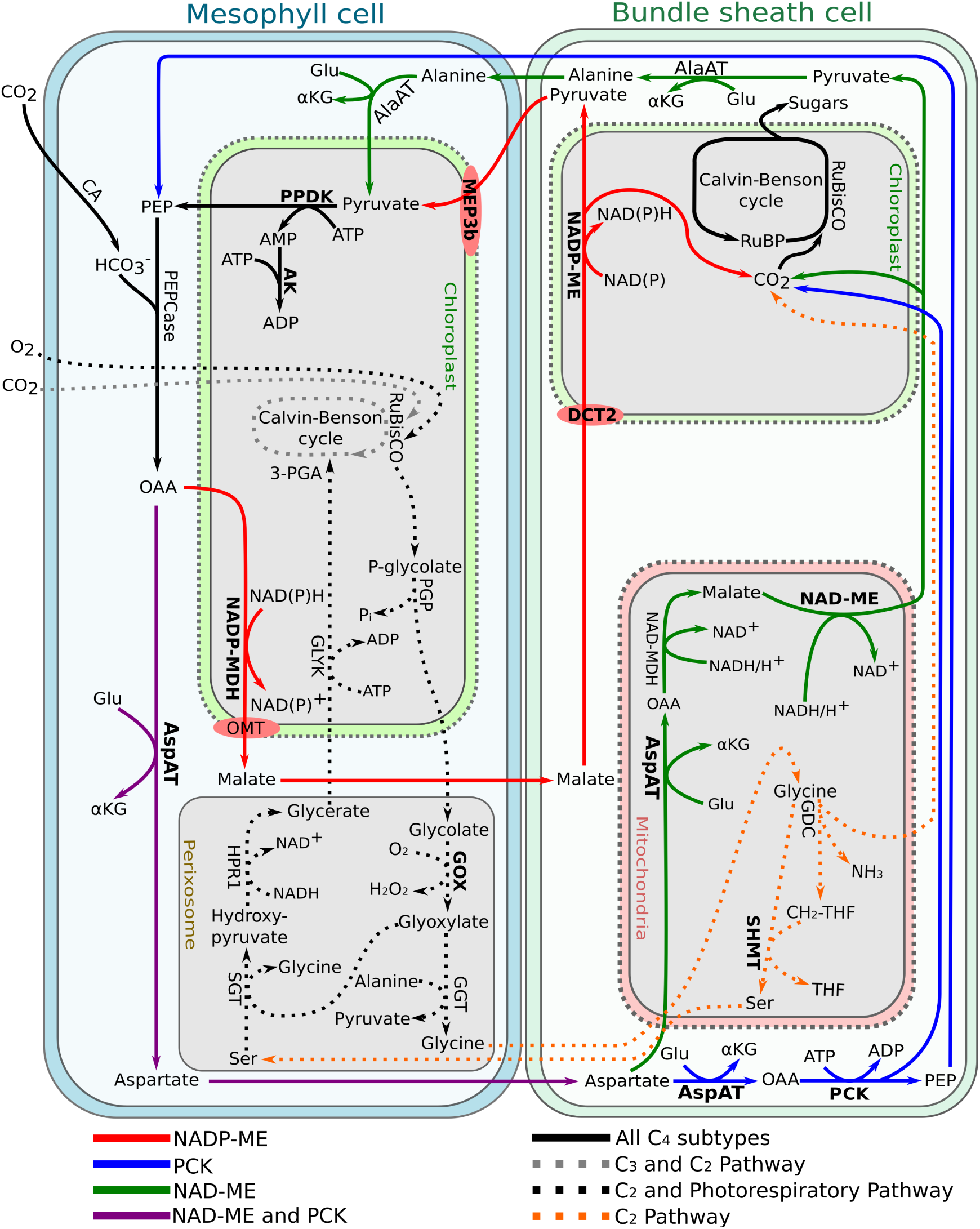
Simplified pathway representation of the three main C_4_ photosynthesis subtypes including the C_3_, C_2_ and photorespiratory pathways. Enzymes studied here are represented in bold. Mitochondrial pathway of the C_2_ cycle is the same as the mitochondrial photorespiratory cycle. However the mitochondrial pathway occurs in the bundle sheath cell in the C_2_ cycle and in the mesophyll cell in the photorespiratory cycle.

## Discussion

The emergence of C_4_ photosynthesis in more than sixty plant lineages remains a fascinating example of the parallel evolutionary emergence of a complex trait. Reconstruction of protein sequence evolution on ancient branches provides a potential window into the mechanisms plants employed to cope with a comparatively rapid drop in CO_2_ concentrations in the atmosphere. Here we used sequence data from eight grass species to examine the patterns of evolution in an ancestral lineage that split from its closest sequenced C_3_ relative 18 million years ago, and gave rise to a diverse set of NADP-ME, NAD-ME and PCK utilizing C_4_ species 13 million years ago (Kumar et al., 2017). What happened over that five million year span? Did that ancestral lineage make the leap from C_3_ photosynthesis to C_4_, or was it a matter of luck or some genetic predisposition that lead to all of its extant descendants utilizing C_4_ photosynthesis, admittedly diverse varieties of C_4_ photosynthesis today?

If one were to focus only on core enzymes which are both low copy and clearly necessary for all forms of the C_4_ photosynthetic pathway, it would appear that this ancient lineage did indeed make the transition from C_3_ photosynthesis to C_4_ (Figure S1). A focus on enzymes used in some C_4_ pathways but not others instead is consistent with the common ancestor of the MPC clade still utilizing C_3_ photosynthesis. This model also implies C_4_ photosynthesis instead emerged later and independently in the diversification of this clade. The pattern we observed is also inconsistent with a clean separation into NADP-ME, NAD-NE, and PCK utilizing species (Figure S2). In fact only in two cases, AspAT and NAD-ME, where the branches that exhibited accelerated protein sequence evolution relative to C_3_ outgroups, are entirely consistent with the primary C_4_ pathways utilizing by species descended from those branches (Table 2). This complexity of C_4_ pathway utilization suggested by which enzymes show accelerated rates of protein sequence evolution in which lineages is consistent with more recent studies. Multiple reports indicate that many species traditionally thought to employ only a single C_4_ pathway may actually fix significant proportions of their total carbon through two or more C_4_ pathways (Walker et al., 1997; John et al., 2014; Huang et al., 2016; de Oliveira Dal’Molin et al., 2016; Washburn et al., 2017).

The observation that glycolate oxidase, a critical component of the photorespiratory pathway, also shows accelerated protein sequence evolution in the lineage leading to the common ancestor of the MPC clade (Figure S3) in addition to PPDK and PPDK-RP suggests an intermediate hypothesis. Sometime in the five million year span between the divergence of the MPC lineage from *Dichanthelium oligosanthes* and the most recent common ancestor of the MPC clade, this lineage transitioned from conventional C_3_ photosynthesis to the intermediate C_2_ photosynthetic cycle where the photorespiratory pathway acts as carbon pump, rather than utilizing any of the three decarboxylation enzymes of classical C_4_ photosynthesis (Figure 4). This model is consistent with (Washburn et al., 2015) where ancestral state reconstruction suggested the MPC species utilizing different subtypes of C_4_ photosynthesis may have evolved from a non-C_4_ ancestor.

It is intriguing to speculate that these ancient changes in protein sequence represents an evolutionary echo of the strong selection acting on plants to adapt to a dramatic decline in atmospheric CO_2_ levels. However, it is important to keep in mind the same caveat discussed above: elevated dN/dS ratios which remain below one, even when statistically significant, can be explained by either a mixture of purifying and positive selection or by a simple relaxation of purifying selection. In addition, parallel substitution for the same amino acid substitutions in sister lineages can sometimes lead to a amino acid change being incorrectly inferred to have happened a single time in the common ancestor. This phenomenon has been observed in the study of the parallel evolution of C_4_ grasses in the past (Christin et al., 2007, 2008). The findings we present here are suggestive of a C_2_ intermediate in the evolution of C_4_ photosynthesis in the MPC clade (Sage, 2001; Edwards, 2019), but they do not yet represent conclusive proof of such an ancestor. Given the accelerating pace of plant genome sequencing and improved automated methods for identifying orthologs even in the absence of high quality synteny data, future investigations may incorporate data from larger numbers of C_3_ and C_4_ grass species, as well as larger numbers of different genes known to be or believed to be involved in C_3_, C_2_, and/or C_4_ photosynthesis.

## Data availability

CDS sequences for *Urochloa fusca* are available at Zenodo with the identifier 10.5281/zenodo.3238541.

## Author Contributions

DSC, SKKR, YZ and JCS wrote the paper; DSC and JCS designed and conducted the analyses; JCS and YZ collected the data.

## Acknowledgements

This study was supported by Science without Borders scholarship (214038/2014-9) to DSC, a Robert B. Daugherty Water for Food Institute research support to JCS and award 2016-67013-24613 from the USDA National Institute of Food and Agriculture to JCS. We thank Jyothi Kumar for critical reading and commentary on earlier versions of this manuscript.

## Supplemental Data

### Supplementary material

File S1) Complete set of supplementary figures

File S2) The syntenic and and recriptocal best blast hit inferred orthologs used in this study

**Figure S1:**
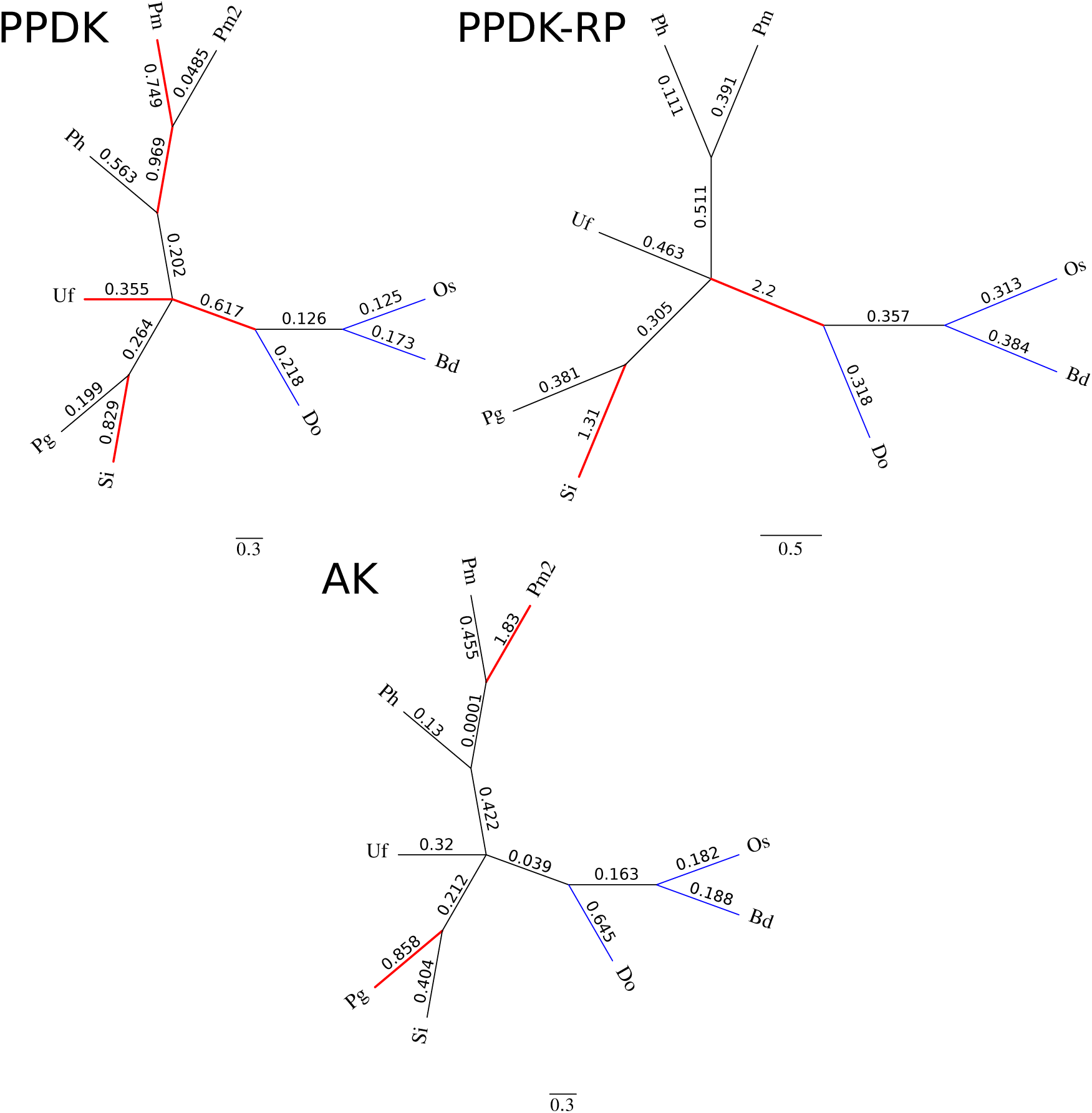
Unrooted phylogenetic trees of C_4_ photosynthesis core enzymes, present in all subtypes, according to citations in Table 1. Branch lengths are equal. Thick red branches represent branches evolving significantly faster than background C_3_ branches in blue. Abbreviations: Os = *Oryza sativa*, Bd = *Brachypodium distachyon*, Do = *Dichanthelium oligosanthes*, Si = *Setaria italica*, Pg = *Pennisetum glaucum*, Uf = *Urochloa fusca*, Ph = *Panicum hallii*, Pm = *Panicum miliaceum*.

**Figure S2:**
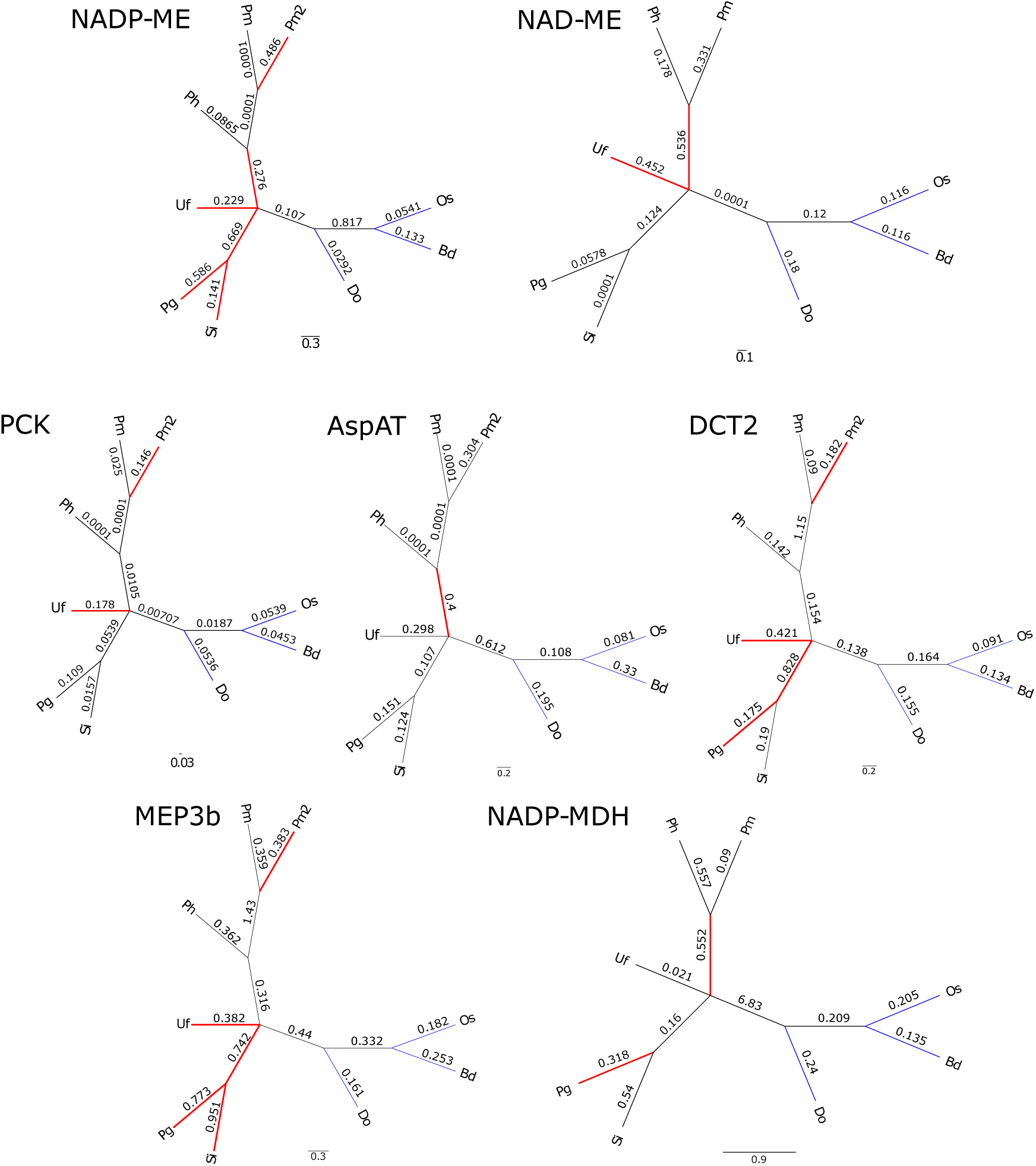
Unrooted phylogenetic trees of C_4_ photosynthesis subtype specific enzymes. Branch lengths are equal. Thick red branches represent branches evolving significantly faster than background C_3_ branches in blue. Abbreviations: Os = *Oryza sativa*, Bd = *Brachypodium distachyon*, Do = *Dichanthelium oligosanthes*, Si = *Setaria italica*, Pg = *Pennisetum glaucum*, Uf = *Urochloa fusca*, Ph = *Panicum hallii*, Pm = *Panicum miliaceum*.

**Figure S3:**
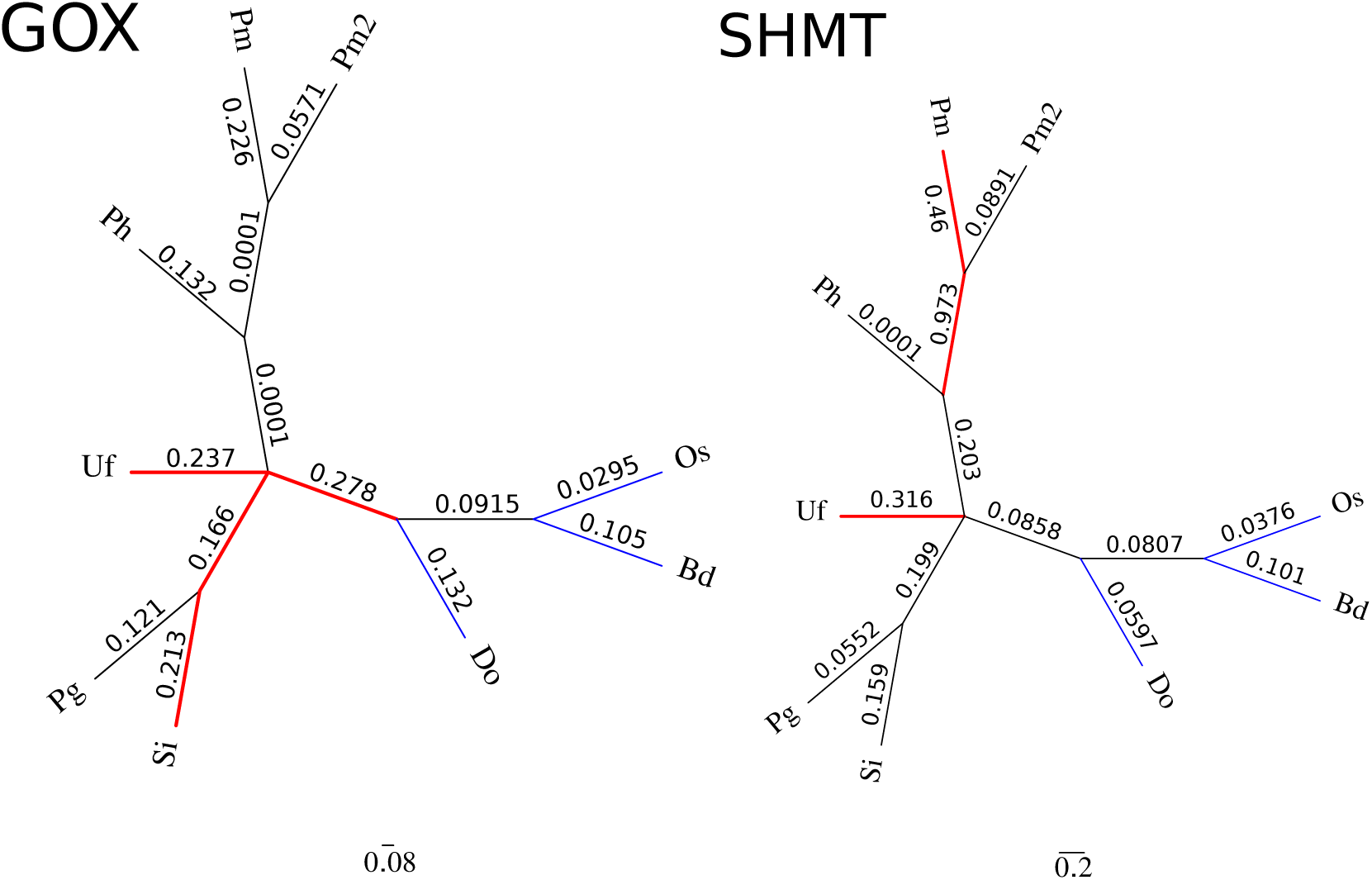
Unrooted phylogenetic trees of C_4_ photosynthesis photorespiratory enzymes. Branch lengths are equal. Thick red branches represent branches evolving significantly faster than background C_3_ branches in blue. Abbreviations: Os = *Oryza sativa*, Bd = *Brachypodium distachyon*, Do = *Dichanthelium oligosanthes*, Si = *Setaria italica*, Pg = *Pennisetum glaucum*, Uf = *Urochloa fusca*, Ph = *Panicum hallii*, Pm = *Panicum miliaceum*.

