## Supplemental Material 1 for "Reconstructed protein sequence evolution consistent with the evolution of C_4_ photosynthesis via a C_2_ ancestor in the Paniceae"

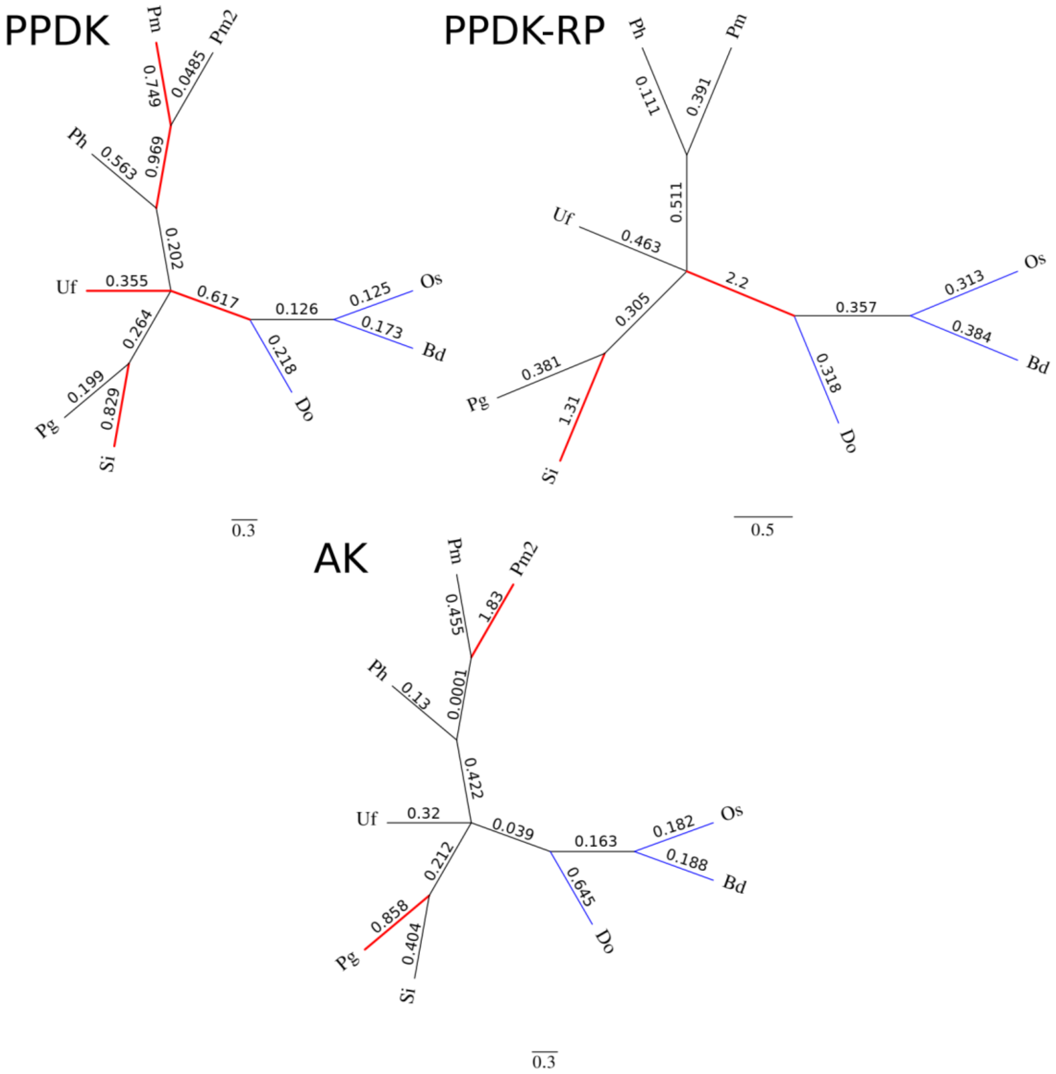


Figure S1: **Unrooted phylogenetic trees of $C_4$ photosynthesis core enzymes, present in all subtypes, according to citations in Table 1.** Branch lengths are equal. Thick red branches represent branches evolving significantly faster than background $C_3$ branches in blue. Abbreviations: Os = *Oryza sativa*, Bd = *Brachypodium distachyon*, Do = *Dichanthelium oligosanthes*, Si = *Setaria italica*, Pg = *Pennisetum glaucum*, Uf = *Urochloa fusca*, Ph = *Panicum hallii*, Pm = *Panicum miliaceum*.

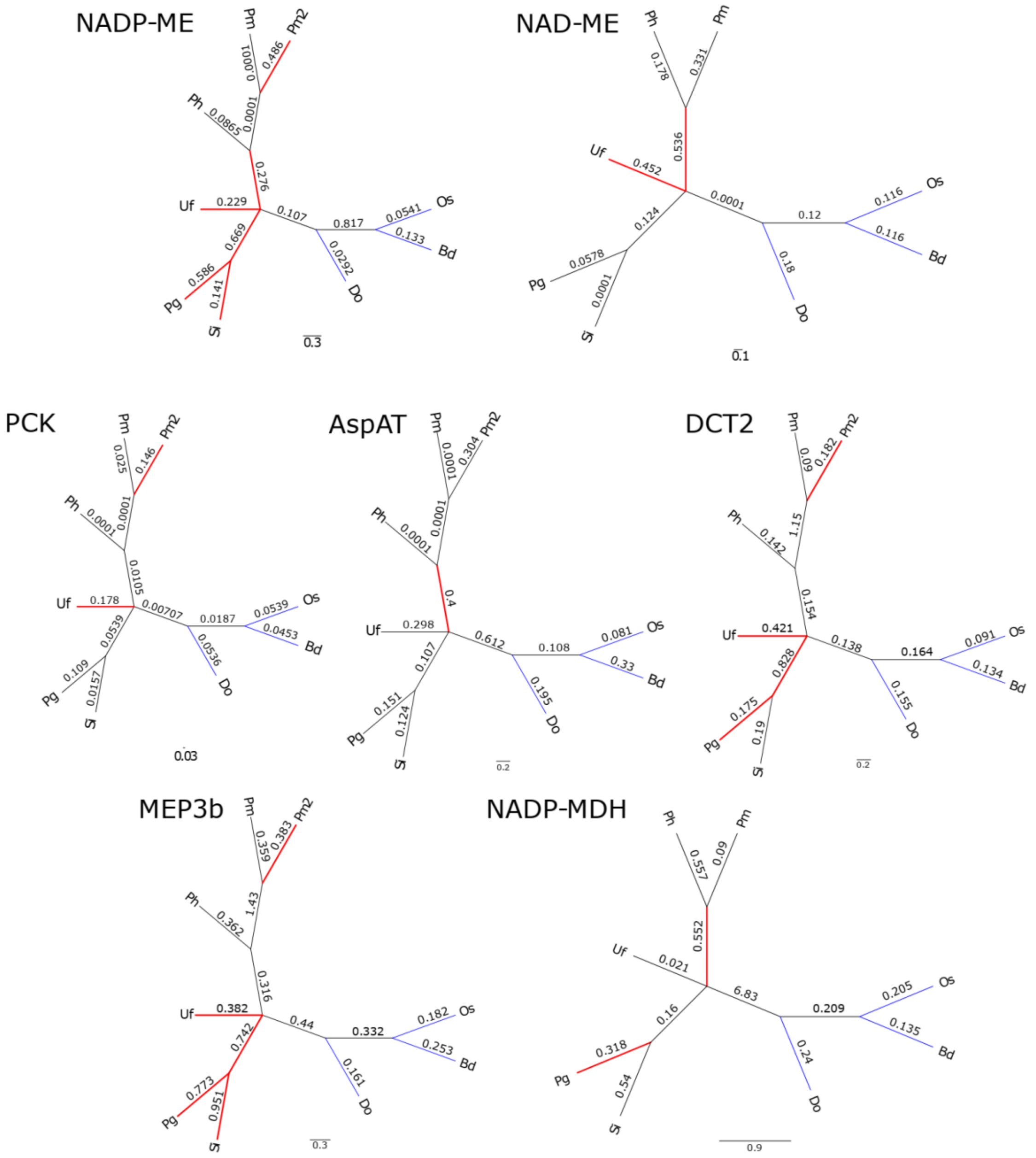


Figure S2: **Unrooted phylogenetic trees of $C_4$ photosynthesis subtype specific enzymes.** Branch lengths are equal. Thick red branches represent branches evolving significantly faster than background $C_3$ branches in blue. Abbreviations: Os = *Oryza sativa*, Bd = *Brachypodium distachyon*, Do = *Dichanthelium oligosanthes*, Si = *Setaria italica*, Pg = *Pennisetum glaucum*, Uf = *Urochloa fusca*, Ph = *Panicum hallii*, Pm = *Panicum miliaceum*.

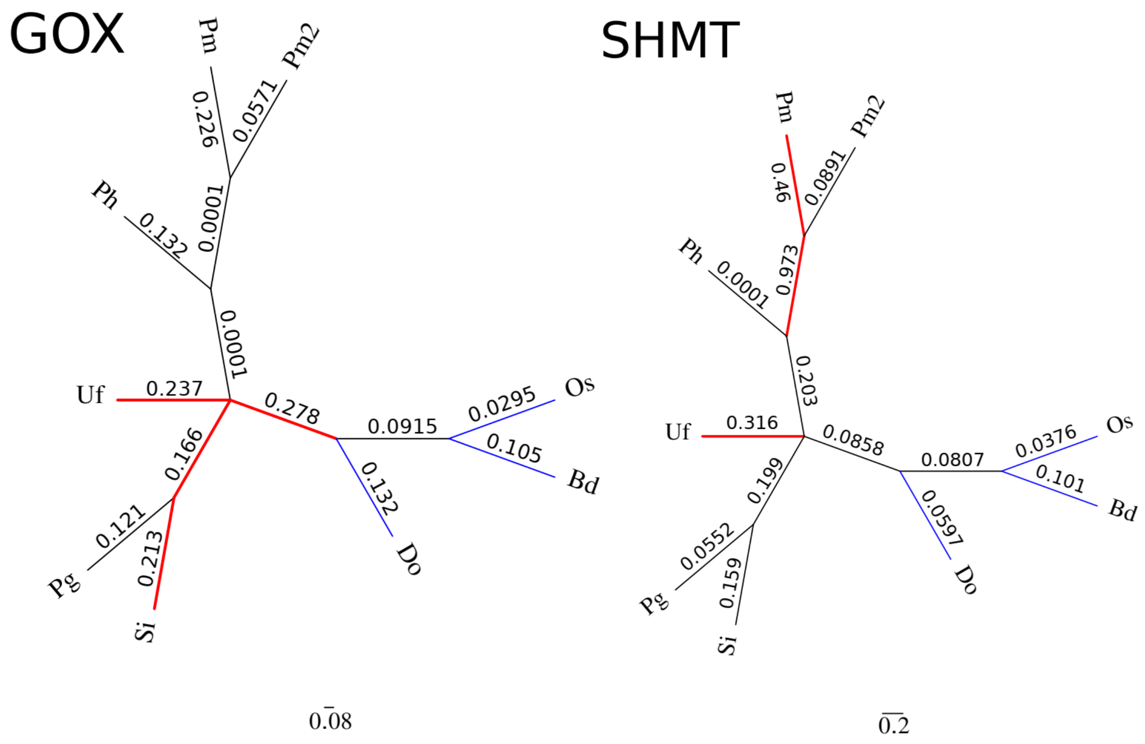


Figure S3: **Unrooted phylogenetic trees of $C_4$ photosynthesis photorespiratory enzymes.** Branch lengths are equal. Thick red branches represent branches evolving significantly faster than background $C_3$ branches in blue. Abbreviations: Os = *Oryza sativa*, Bd = *Brachypodium distachyon*, Do = *Dichanthelium oligosanthes*, Si = *Setaria italica*, Pg = *Pennisetum glaucum*, Uf = *Urochloa fusca*, Ph = *Panicum hallii*, Pm = *Panicum miliaceum*.
